# Targeted proteomic quantitation of NRF2 signaling and predictive biomarkers in HNSCC

**DOI:** 10.1101/2023.03.13.532474

**Authors:** Nathan T. Wamsley, Emily M. Wilkerson, Li Guan, Kyle M. LaPak, Travis P. Schrank, Brittany J. Holmes, Robert W. Sprung, Petra Erdmann Gilmore, Sophie P. Gerndt, Ryan S. Jackson, Randal Paniello, Patrik Pipkorn, Sidarth V. Puram, Jason T. Rich, Reid R. Townsend, José P. Zevallos, Paul Zolkind, Quynh-Thu Le, Dennis Goldfarb, M. Ben Major

**Author notes:** **Corresponding Authors** Dr. Michael B. Major, Washington University in St. Louis, Department of Cell Biology & Physiology, St. Louis, MO, 63110, USA, Dr. Dennis Goldfarb, Washington University in St. Louis, Department of Cell Biology & Physiology, St. Louis, MO, 63110, USA.

## Abstract

The NFE2L2/NRF2 oncogene and transcription factor drives a gene expression program that promotes cancer progression, metabolic reprogramming, immune evasion and chemoradiation resistance. Patient stratification by NRF2 activity may guide treatment decisions to improve outcome. Here, we developed a mass spectrometry (MS)-based targeted proteomics assay based on internal standard triggered parallel reaction monitoring (IS-PRM) to quantify 69 NRF2 pathway components and targets as well as 21 proteins of broad clinical significance in head and neck squamous cell carcinoma (HNSCC). We improved the existing IS-PRM acquisition algorithm, called SureQuant^™^, to increase throughput, sensitivity, and precision. Testing the optimized platform on 27 lung and upper aerodigestive cancer cell models revealed 35 NRF2 responsive proteins. In formalin-fixed paraffin-embedded (FFPE) HNSCCs, NRF2 signaling intensity positively correlated with NRF2 activating mutations and with SOX2 protein expression. PD-L2/CD273 and protein markers of T-cell infiltration correlated positively with one another and with human papilloma virus (HPV) infection status. p16/CDKN2A protein expression positively correlated with the HPV oncogenic E7 protein, and confirmed the presence of translationally active virus. This work establishes a clinically actionable HNSCC protein biomarker assay capable of quantifying over 600 peptides from frozen or FFPE archived tissues in under 90 minutes.

## Introduction

Head and neck squamous cell carcinoma (HNSCC) is the seventh most common cancer worldwide; in the United States, 66,000 new cases and 15,000 deaths were expected in 2022 [1, 2]. Key risk factors include alcohol consumption, tobacco use, and human papilloma virus (HPV) infection [1]. Nearly all HPV(+) tumors of the head and neck affect the oropharynx and are associated with a favorable prognosis of a 75-80% 5-year survival rate [3, 4]. Immunohistochemistry (IHC) staining for p16/CDKN2A serves as a proxy for HPV status and is the leading prognostic biomarker in HNSCC [3]. In contrast, HPV(-) oropharyngeal cancer carries a 5-year survival rate of 45–50% [4]. For locoregionally advanced disease, radiation therapy (RT) or surgery with RT has remained the first line treatment for decades, with no meaningful improvement [1, 5]. The recent inclusion of immune checkpoint inhibitors (ICI) to the HNSCC therapeutic armament elicits favorable responses, but only for 20% of patients [6]. Molecular characterization of therapeutic responsive, non-responsive, and recurrent HPV(-) HNSCC have revealed several key determinants of patient outcome. Principal among these is the NRF2 oxidative stress response pathway. Mutations drive constitutive NRF2 activation in 17% of HNSCCs [7]. Active NRF2 signaling in HNSCC prognosticates poor overall survival and predicts locoregional failure following RT [8–14]. However, clinical assays that leverage NRF2 activity to stratify patients for improved therapeutic response remain to be developed and proven. Similarly, current molecular diagnostics fail to identify a majority of ICI non-responders [15]. Here we developed a targeted proteomics assay for HNSCC that quantifies markers for: 1) HPV infection, 2) NRF2 signal transduction, 3) HNSCC tumor suppressors and oncogenes, and 4) immuno-oncology markers.

Constitutive NRF2 signaling drives RT resistance and locoregional failure in HPV(-) HNSCC. In normal cells the KEAP1/CUL3 E3 ubiquitin ligase complex binds NRF2 and catalyzes its ubiquitylation for subsequent proteasomal degradation [16–18]. Metabolic, oxidative, and electrophilic stressors inhibit NRF2 degradation by KEAP1/CUL3, resulting in NRF2 stabilization, nuclear translocation and transcriptional activation of target genes that collectively restore cell health [19]. These targets encode enzymes supporting antioxidant pathways, drug metabolizing enzymes, components of the pentose phosphate pathway, and others [7, 19]. We recently showed that NRF2 activating mutations predict locoregional failure to RT in oral and larynx cancers [13, 14]. However, these mutation-based studies fail to account for many NRF2 active cancers that lack a known mutational driver [7]. A robust, fast, and cost-effective NRF2 activity diagnostic will guide patient treatment decisions, including radiation dose or surgery [20, 21].

Effective biomarkers might also accurately predict successful immune checkpoint inhibition, a therapy to which fewer than one-in-five HNSCC patients respond [15, 22, 23]. Two widely studied prognostic indicators are PD-L1 expression and T-cell inflamed gene expression profile (GEP) [15]. IHC staining for PD-L1 fails to reliably predict ICI response [24]. GEP can identify subsets of patients least likely to respond but only at false positivity rates well exceeding 50% [23, 25, 26]. Immunostaining assays are confounded by poor correspondence of IHC scores to molar abundance, covalent protein modifications and functional redundancy (eg. PD-L2). Further, GEP considers mRNAs that may correlate poorly with their protein counterparts [27-30]. Lastly, a growing body of evidence suggests that an active NRF2 pathway reduces the strength of anti-tumor immunity. In the context of an inflamed tumor microenvironment (TME), NRF2 promotes PD-L1 expression, the recruitment of immunosuppressive myeloid cells, and M2 macrophage polarization [14, 23, 31–33]. A MS-based proteomics tool to quantify both NRF2 signaling and the presence and state of immune cells in the TME might improve prediction of therapeutic response and empower future studies of an NRF2-immune infiltration axis in cancer [23, 31–35].

This work presents an optimized proteomics assay for the study of biomarkers for HPV infection, NRF2 signaling proteins, T-cell infiltration and HNSCC-associated cancer drivers. The technology is based on a custom implementation of internal standard-triggered parallel reaction monitoring, which leverages stable isotope labeled (SIL) peptides to direct efficient data acquisition [36]. IS-PRM enables relative and absolute quantitation of many hundreds of analytes at low-attomolar abundance from minimal sample input, which is suitable to quantify many transcription factors, kinases and other scarce signaling molecules from tumor biopsies and archival tissue blocks [36-39]. We benchmarked our optimized IS-PRM (OIS-PRM) method against a commercial implementation called SureQuant^™^ to establish its improved performance [37, 40]. We applied OIS-PRM to study NRF2 signaling components and targets in genetically engineered cell models and a cohort of genotyped lung, esophageal and head and neck cancer cell lines. Additionally, in two patient cohorts of formalin-fixed paraffin embedded (FFPE) HNSCC tumors, we quantified protein expression for tumor-immune infiltration, pan-squamous cell carcinoma cancer drivers, HPV infection and NRF2 related proteins.

## Results

### Benchmarking an optimized IS-PRM method

Two targeted proteomics methods, IS-PRM and its commercial implementation called SureQuant^™^, achieve sensitive and reproducible quantification of peptides by monitoring spiked-in SIL peptides to direct efficient data acquisition. These internal standard peptides co-elute with their endogenous counterparts so that IS-PRM triggers time-intensive, quantitative “quant” scans targeting an endogenous peptide. Quant scans of the endogenous peptide are triggered by fast “watch” scans which detect and identify the highly-abundant SIL peptide. As a consequence of this efficient use of instrument time, IS-PRM enables quantification of more proteins in a single analysis than a standard PRM method (Fig. 1A; see Supplementary Text). However, through our research with SureQuant^™^, we observed that inefficient scan scheduling on Tribrid^™^ mass spectrometers resulted in excess Orbitrap idle time (Fig. 1B). Using the Thermo Scientific^™^ Tribrid^™^ IAPI, we implemented an optimized IS-PRM method (OIS-PRM) that postpones quant scans until the completion of all SIL-detection watch scans (Fig. 1B).

**Figure 1.**
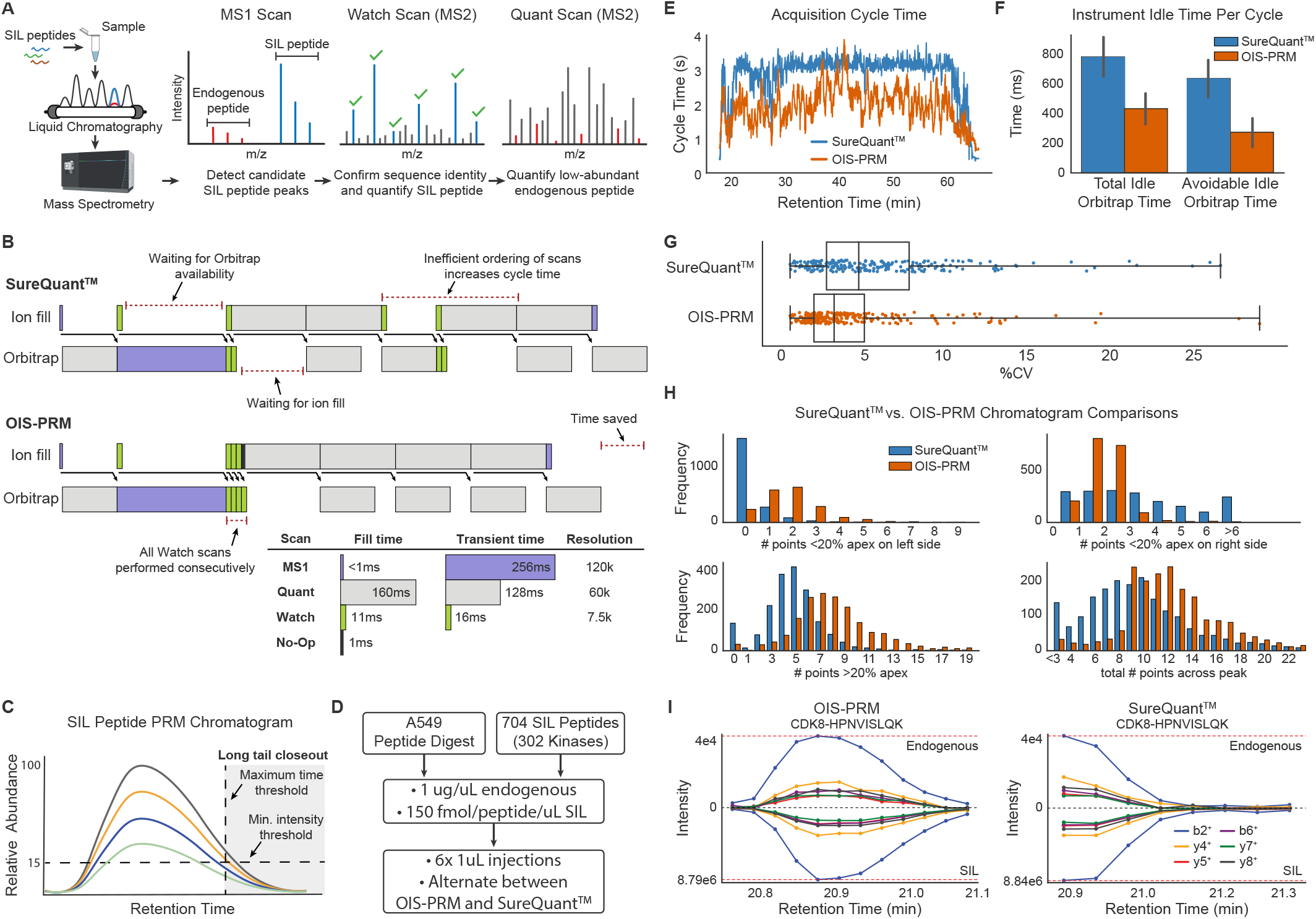
OIS-PRM enabled efficient targeted proteomic data acquisition. **A)** Schematic of the SureQuant^™^ (Thermo Scientific) IS-PRM acquisition algorithm. Stable isotope labeled (SIL) peptides corresponding to each endogenous targeted peptide are spiked into each sample. The heavy SIL peptides are highly abundant, direct efficient data acquisition of endogenous light peptides, and serve as internal standards for quantification. MS1 detection of a SIL peptide triggers an MS2 “Watch” scan for the SIL peptide. When a “Watch” scan confirms a SIL peptide by the presence of pre-determined fragment ions, SureQuant^™^ triggers a “Quant” MS2 scan targeting the endogenous counterpart. Signal intensity from each endogenous peptide is normalized to its corresponding SIL internal standard peptide. **B)** Example of instrument utilization on a Tribrid^™^ mass spectrometer using SureQuant^™^ or our optimized internal standard PRM (OIS-PRM) method. OIS-PRM contiguously performs all “Watch” and “Quant” scans. **C)** Illustration of thresholds used to stop data acquisition of long elution tails. “Watch” and “Quant” scans are not acquired on a target after the summed fragment ion intensity for the SIL peptide falls below a proportion of the maximum observed for that SIL peptide, or after a maximum time since its initial detection. **D)** Experimental design for the comparison of SureQuant^™^ and OIS-PRM methods in the subsequent panels. **E)** Cycle time plot for representative injections. SureQuant^™^ cycle times were capped at a maximum of 3 seconds. **F)** Comparison of Orbitrap idle time. Avoidable idle time subtracts the transient of the MS1 scan from the total. **G)** Comparison of coefficients of variation (CV) for endogenous peptides over three replicate injections. **H)** Number of peptides whose quantified chromatograms satisfied the indicated constraints for all three replicate injections. **I)** Chromatograms for the same peptide in back-to-back injections using OIS-PRM and SureQuant^™^.

Additionally, we hypothesized that avoiding quant scans during the long tail of a peptide elution profile would free up instrument time without compromising quantitative accuracy; therefore, we added thresholds for minimum relative intensity and maximum elution time (Fig. 1C). Last, we reordered quant scans to prioritize newly detected peptides and capture the starts of their elution profiles.

To evaluate OIS-PRM performance, we ran six injections of an A549 lung cancer cell line digest, alternating between SureQuant^™^ and OIS-PRM (Fig. 1D). The SIL library included 704 kinase-associated peptides analyzed with a 50min liquid chromatography (LC) gradient. OIS-PRM reduced the median cycle time from 3.1s to 1.8s (Fig. 1E), and efficient scan ordering contributed 360ms per cycle to this difference (Fig. 1F). With maximum time and intensity thresholds, the number of SIL peptides with at least 7 points across the peak improved from 173 to 252, and median CV from 4.6 to 3.2 percent for OIS-PRM compared to SureQuant^™^ (Fig. 1G). Overall, OIS-PRM and SureQuant^™^ quantified 264 and 259 peptides, respectively, with CVs less than 20%. OIS-PRM sampled more points per peak, missed fewer peak fronts and oversampled from peak tails less often (Fig. 1H-I). Finally, we compared OIS-PRM to a data-dependent acquisition method (DDA). Over three replicate injections, the OIS-PRM method quantified 172 out of 302 kinases with a CV less than 20% (Supplementary Fig. 1). Although DDA identified 4680 protein groups on average, only 47 kinases were quantified with a CV under 20%.

### Development of an Internal Standard Peptide Array for HNSCC

Aberrant NRF2 activity prognosticates resistance to radiation and chemotherapy in cancers of the lung and upper airway. Therefore, to empower the clinical potential of OIS-PRM, we developed an NRF2 and HNSCC-specific SIL catalogue. This resource includes 227 peptides that represent 90 proteins: 68 NRF2 interacting proteins or transcriptional targets; 10 immuno-oncology markers that include immune checkpoint proteins, cytokines, T-cell surface markers, and immuno-oncology markers; 8 known SCC tumor suppressors and oncogenes; HPV E6 and E7, GAPDH, and DHFR. To develop the NRF2-activity SIL panel, we began with 23 well-established NRF2 targets and NRF2 interacting proteins. For expansion, we proteogenomically analyzed the Clinical Proteomic Tumor Analysis Consortium (CPTAC) LUAD, LUSC, and HPV(-) HNSCC cohorts, which frequently contain NRF2 pathway activating mutations (Supplementary Fig. 2) [41–43]. We sorted the tumors by the abundance of the 13 NRF2 target proteins that were expressed in all of the tumors (Fig. 2A). Principal component analysis compressed these data into a single “NRF2 activity score” that captured 60% of the data variance (Fig. 2B). Overall, NRF2/KEAP1 mutated tumors reported higher NRF2 activity scores than tumors lacking mutations (Fig. 2C). However, many tumors with mutations showed low expression of NRF2 target genes and conversely, many wild-type tumors overexpressed NRF2 targets. It follows that an assay to classify tumors as NRF2 active based on genotype alone would give many false positives and false negatives (Fig. 2C).

**Figure 2.**
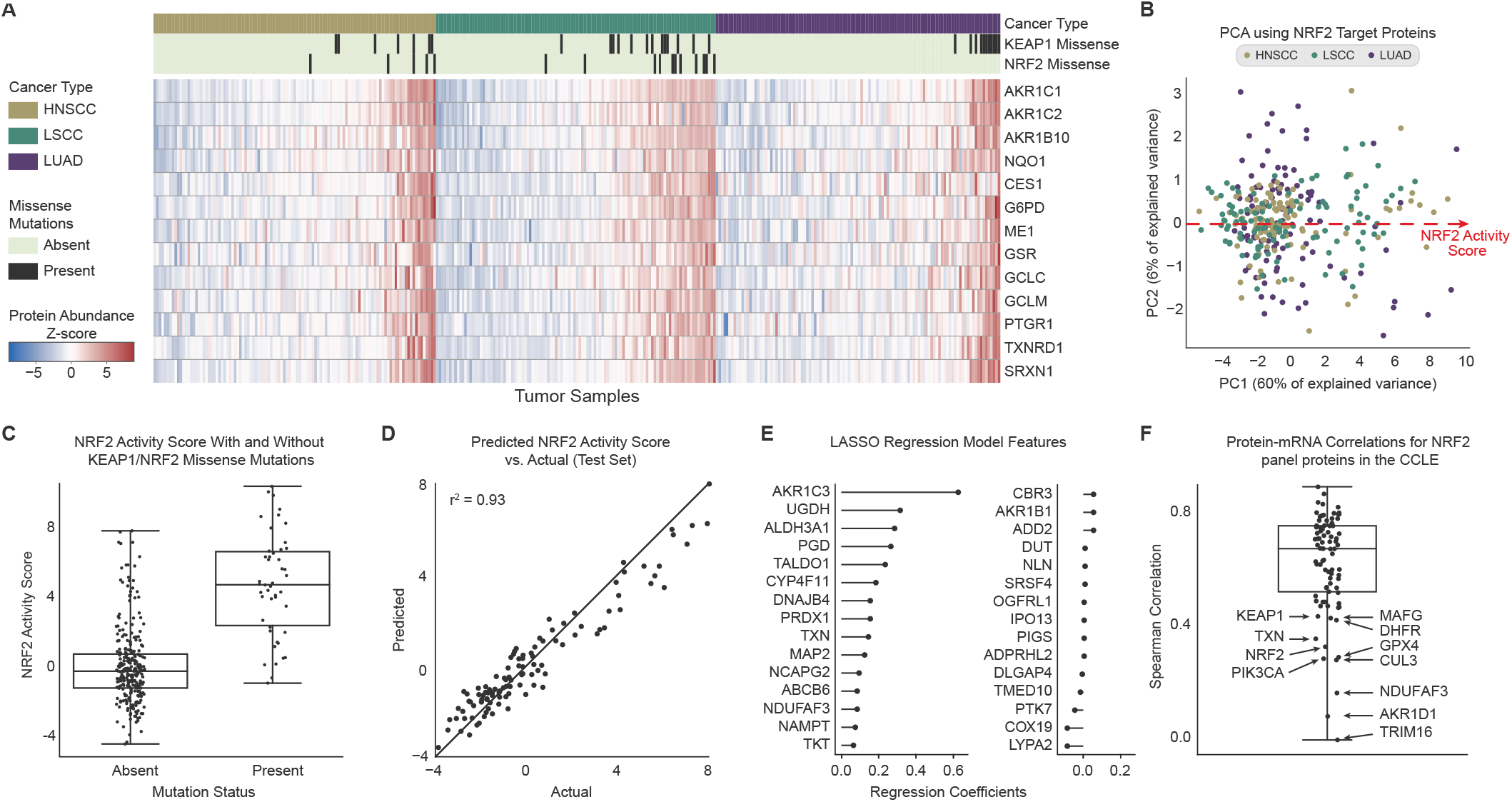
Combined analysis of three Clinical Proteomics Tumor Analysis Consortium cohorts showed that a strongly correlated subset of proteins could predict NRF2/KEAP1 genotype. **A)** Protein expression of thirteen NRF2 targets in CPTAC cohorts for HNSCC, LSCC, and LUAD. NRF2 mutations distant from the ‘DLG’ or ‘ETGE’ motifs were excluded from the count. **B)** PCA plot of the first two principal components for the CPTAC data by the thirteen NRF2 target peptides. The NRF2 activity score refers to the position along the 1st principal component of the data. **C)** Distribution of NRF2 scores for the combined CPTAC cohorts split based on the presence or absence of missense mutations in KEAP1 or NRF2. **D)** A LASSO regression model trained to predict the NRF2 scores for tumors given the 6133 proteins not included in the original set of thirteen NRF2 targets. The model was trained by 10-fold cross validation on a training set of 2/3 of the data (218 tumors). Predicted NRF2 scores are plotted against the true NRF2 pathway scores for an independent test set containing one-third of the data (109 tumors). **E)** Feature weights for the regression model in **C. F)** Protein to mRNA correlations for the NRF2 IS-PRM panel proteins across the cancer cell line encyclopedia.

We therefore used the CPTAC data to identify additional proteins useful for monitoring the NRF2 pathway. A LASSO regression was trained on >6000 proteins not among the initial 13 NRF2 targets to predict the NRF2 activity scores across the CPTAC cohorts (Fig. 2D). We found that a mere 30 proteins with non-zero coefficients in the model could accurately predict the NRF2 scores (Fig. 2E). Literature evidence for most of those proteins with non-negligible coefficients supported their status as NRF2 targets. We included 17 of these in the final SIL peptide array (Supplementary Tables 1–2). Finally, we inspected the mRNA to protein correlations within our NRF2 panel using the Cancer Cell Line Encyclopedia [30]. While most NRF2 transcriptional targets encode enzymes that correlated well, key regulatory proteins NRF2, KEAP1, MAFG, CUL3, and TRIM16 correlated poorly (Fig. 2F). Similarly, several immune checkpoint proteins correlated poorly with their mRNA abundances, which agrees with recent reports [28–30, 44].

### Validation of the NRF2 pathway in cell-lines

To assess whether our IS-PRM assay could quantify NRF2 pathway activation, we engineered HPV(+) HNSCC cell lines SCC90, SCC152, and SCC154 to stably express NRF2^E79Q^, a common cancer-associated activating NRF2 mutation (Fig. 3A) [45]. Stable expression of NRF2^E79Q^ protein in each cell line induced the expression of NRF2 and two of its canonical targets, NQO1 and HMOX1 (Fig. 3B). IS-PRM analyses of the six cell lines revealed that NRF2 activation induced target expression over parental controls between less than 2-fold for some proteins to as high as 90-fold for AKR1B10 (Fig. 3C).

**Figure 3.**
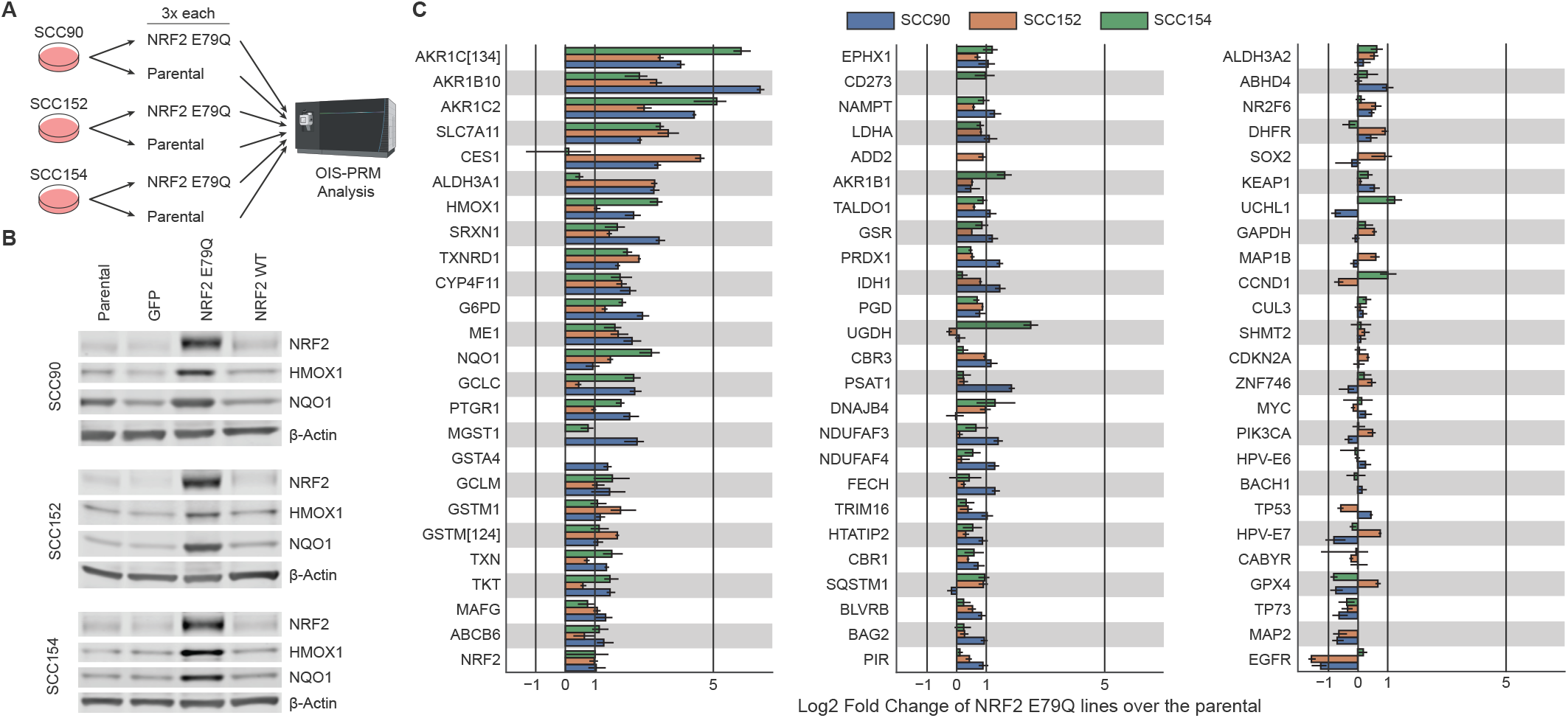
Oral squamous cell carcinoma cell lines that stably overexpressed a cancer derived NRF2 variant likewise overexpressed NRF2 targets at the protein level. **A)** Parental oral squamous cell carcinoma cell lines stably expressing WT NRF2and derived cell lines stably expressing NRF2 E79Q were grown in three replicates each, harvested on the same-day, and analyzed by OIS-PRM **B)** Protein expression measured by Western blot from whole-cell lysates of cell lines derived from human oral squamous cell carcinomas. Blots include parental cell lines and those stably expressing GFP, NRF2 E79Q, and NRF2 WT. **C)** Parental and NRF2 E79Q cell lines as analyzed by OIS-PRM. Error bars represent the minimum and maximum log2 fold change between pairs of parental and NRF2 E79Q replicates.

For additional testing, we applied the IS-PRM method to a collection of 21 cell lines with known *NRF2/KEAP1* genotype and activity status (Fig. 4A). Hierarchical clustering placed all cell lines with *NRF2* or *KEAP1* mutations in the same cluster (Fig. 4B). In agreement with the CPTAC cohorts (Fig. 2A), several cell lines lacking a causative mutation overexpressed NRF2 targets. We modeled expression of NRF2 target proteins using a hierarchical Bayesian analysis (see supplementary methods) and found that despite their wide-spread use as NRF2-activity markers, HMOX1 and NQO1 fell among the less overexpressed NRF2 targets. Even the abundance of NRF2 itself did not perfectly discriminate between the active and inactive cell lines. The posterior density of the logarithmic fold change parameter for GAPDH concentrated around zero and suggested good data normalization.

**Figure 4.**
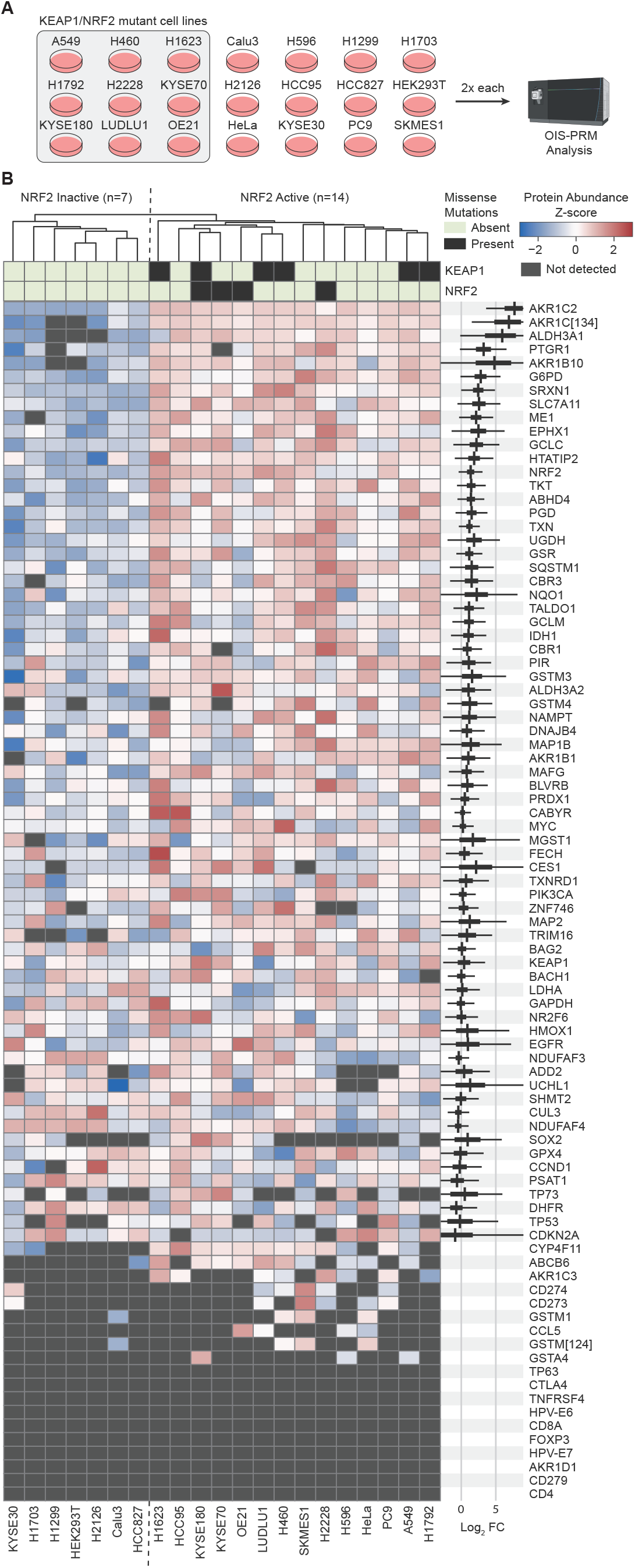
OIS-PRM analysis of 21 cancer cell lines distinguished between NRF2 active and inactive cell lines. **A)** Twenty-one cancer cell lines were cultured in biological duplicate, harvested, and subject to OIS-PRM using a HNSCC-specific SIL peptide array. **B)** Results of the experiment described in **A**. Protein abundances were averaged for each replicate and then data were row normalized by Z-scores and hierarchically clustered. On the right-hand-side the thick black bands contain 95% of the posterior density of the mean logarithmic fold change between the active and inactive clusters. The narrow grey bands contain 95% of the posterior predictive density for the logarithmic fold change in expression between an NRF2 active over an NRF2 inactive cell line.

### OIS-PRM analysis of Head and Neck Squamous Cell Carcinomas

After establishing OIS-PRM in cultured cell models, we next tested it across two sets of archived FFPE HNSCC tumors: 1) 10 HPV(+) and 20 HPV(-) oropharyngeal squamous cell carcinomas collected as 50μm curls (Fig. 5); and 2), punch biopsies from 30 HPV(-) oral squamous cell carcinomas including 20 NRF2 WT tumors and 10 tumors with NRF2^E79Q^ or NRF2^E79K^ activating mutations (Fig. 6). After testing and optimizing a protocol for protein extraction from FFPE, we evaluated protein quality by DDA-MS on 10 HPV(+) oropharyngeal squamous cell carcinomas FFPE curls (Fig. 5A, B). On average, each 50μm curl yielded 300 ug of protein and 18,200 peptides mapping to 3,600 protein groups (Fig. 5B). These yields and overall peptide characteristics were similar to those of prior FFPE proteomic studies [46–49].

**Figure 5.**
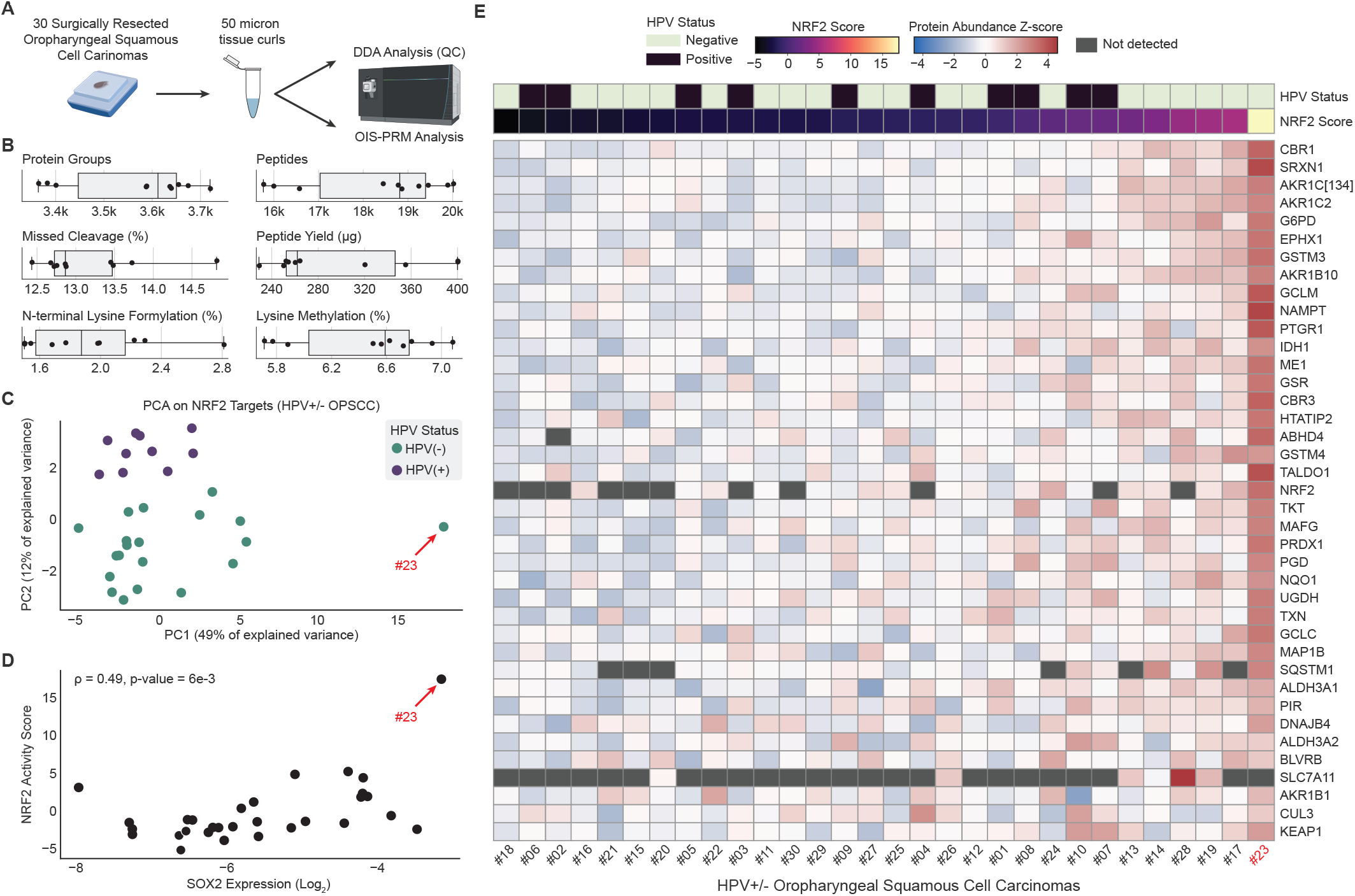
OIS-PRM analysis of FFPE oropharyngeal squamous cell carcinomas revealed a large dynamic range of NRF2 target expression. **A)** Schema describing the collection and analysis of 20 HPV(-) and 10 HPV(+) oropharynx tumors. Protein was extracted from 50-micron curls cut from FFPE tumor blocks and subject to OIS-PRM. **B)** Summary statistics for data-dependent acquisition proteomics on the 10 HPV(+) tumors Lysine methylation and N-terminal lysine formylation are common artifacts of formalin fixation. **C)** PCA plot from protein abundances of differently expressed NRF2 target proteins as measured by OIS-PRM.**D)** Scatterplot of NRF2 activity scores and SOX2 protein abundance for each tumor. The NRF2 score refers to the position along the first principal component from **C. E)** Heatmap of OIS-PRM data with row normalized Z-scores.

**Figure 6.**
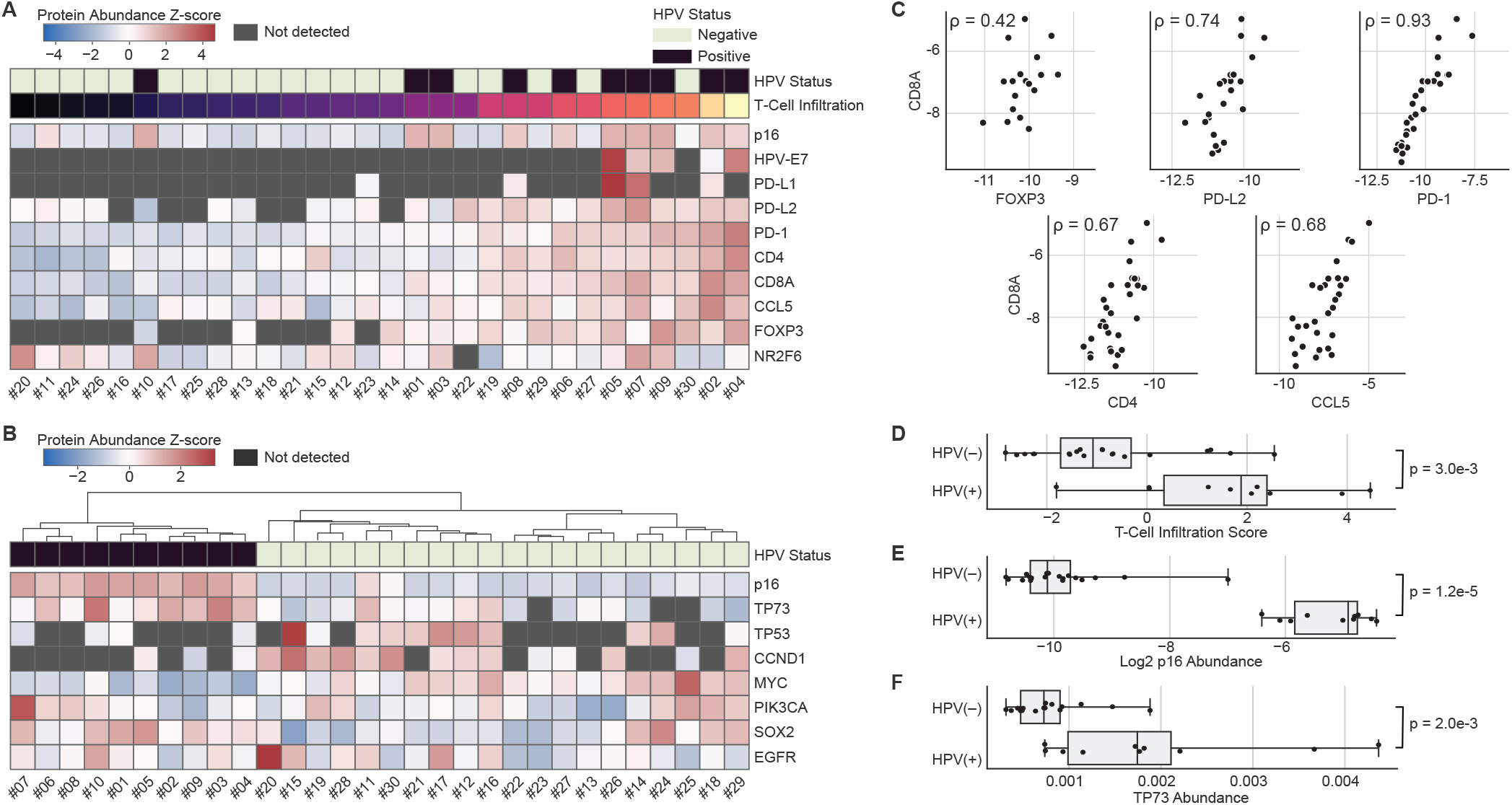
OIS-PRM analysis of FFPE oropharyngeal squamous cell carcinomas reveals HPV-specific differences in the expression of T-cell markers and oncogenes. **A-B)** Expression of immuno-oncology markers and cancer drivers by OIS-PRM for the same oropharynx tumors as in **Figure 5. C)** Spearman correlations between the immuno-oncology markers shown in **A. D-F)** Immune infiltration score, p16 abundance, and TP73 abundance split by HPV status. A PCA analysis of the tumors in **A** using PD-1, CD4, CD8α, CCL5, and FOXP3 expression as features was used to calculate the T-cell infiltration scores as the position of each tumor along the first principal component. P-values were calculated by a two-sided Mann-Whitney U-test.

OIS-PRM analysis of the 30 FFPE curls from HPV(+) and HPV(-) oropharyngeal squamous cell carcinomas revealed expected and novel protein correlations. As with the CPTAC cohort, the first principal component served as an NRF2 score, and it explained fifty percent of the variance (Fig. 5C). This NRF2 score positively correlated with SOX2 protein abundance (Spearman r = 0.49, p-value = 0.006, Fig. 5D). Unexpectedly, we also observed that the second principal component NRF2 score perfectly separated the HPV(+) from the HPV(-) tumors. One of the thirty tumors substantially overexpressed NRF2 targets relative to the others; possibly indicating the presence of a NRF2 activating mutation (Fig. 2C, E).

In the same experiment we quantified immuno-oncology biomarkers and cancer drivers (Fig. 6A-B). The abundances of T-cell associated proteins, CD8α, FOXP3, PD-L2, PD-1, CD4, and CCL5 correlated with one another across the cohort (Fig. 6C). Notably, PD-1 had a near perfect rank correlation with the cytotoxic T-cell maker, CD8α, and with the exception of FOXP3 these rank correlations were as strong or stronger than for the typical protein and its mRNA in the CCLE [30]. Using a principal component analysis, we derived a T-cell infiltration score and found that HPV(+) tumors displayed significantly higher T-cell infiltration than HPV(-) tumors (Fig. 6D). PD-L1 and PD-L2 were detected in 16% and 80% of the tumors, respectively. Nearly all tumors expressed the transcriptional repressor, NR2F6, at detectable levels.

Protein expression of p16/CDKN2A is a commonly used surrogate for HPV infection; direct MS-based detection of HPV has not previously been established. OIS-PRM detected the E6 and E7 proteins in the HPV(+) oral squamous cell carcinoma cell lines, SCC90, SCC152, and SCC154 (Fig. 3C). Across the oropharyngeal squamous cell carcinomas tumor cohort, we detected E7 in five of the HPV(+) tumors and in none of the HPV(-) tumors (Fig. 6A). As expected, p16 expression separated HPV(+) from HPV(-) tumors (Fig. 6E). HPV(+) tumors also significantly overexpressed TP73 compared to HPV(-) tumors, which agrees with a previous report (Fig. 6F) [50].

In addition to the HPV(+) and HPV(-) oropharyngeal squamous cell carcinomas, we also tested OIS-PRM on FFPE tumor punches from a cohort of 19 HPV(-) oral squamous cell carcinomas that were genotyped for NRF2 (Fig. 7A). These included 8 *NRF2*^*E79Q*^ or *NRF2*^*E79K*^ activating/mutant tumors and 11 NRF2 wildtype tumors. Of the 8 NRF2 mutant tumors, 6 strongly expressed NRF2 target genes. Several NRF2-target proteins were expressed at greater than 4-fold in NRF2 mutant compared to wildtype, including NQO1, AKR1C2, GSTM3, GSTM4 and ALDH3A1 (Fig. 7B-C), but some of these proteins were not among the most overexpressed proteins in our cell lines analysis (Fig. 4B). Examining immune cell markers, T-cell infiltration and NRF2 activity did not correlate (Fig. 7D), and likewise, PD-L1 abundance did not correlate with that of PD-L2 (Fig. 7E). Finally, similar to the HPV cohort of oropharyngeal squamous cell carcinomas, SOX2 abundance and NRF2 activity trended to a positive correlation (Fig. 7F). This correlation supports a prior report showing that NRF2 activation associates with SOX2 amplification in squamous cell carcinomas [32].

**Figure 7.**
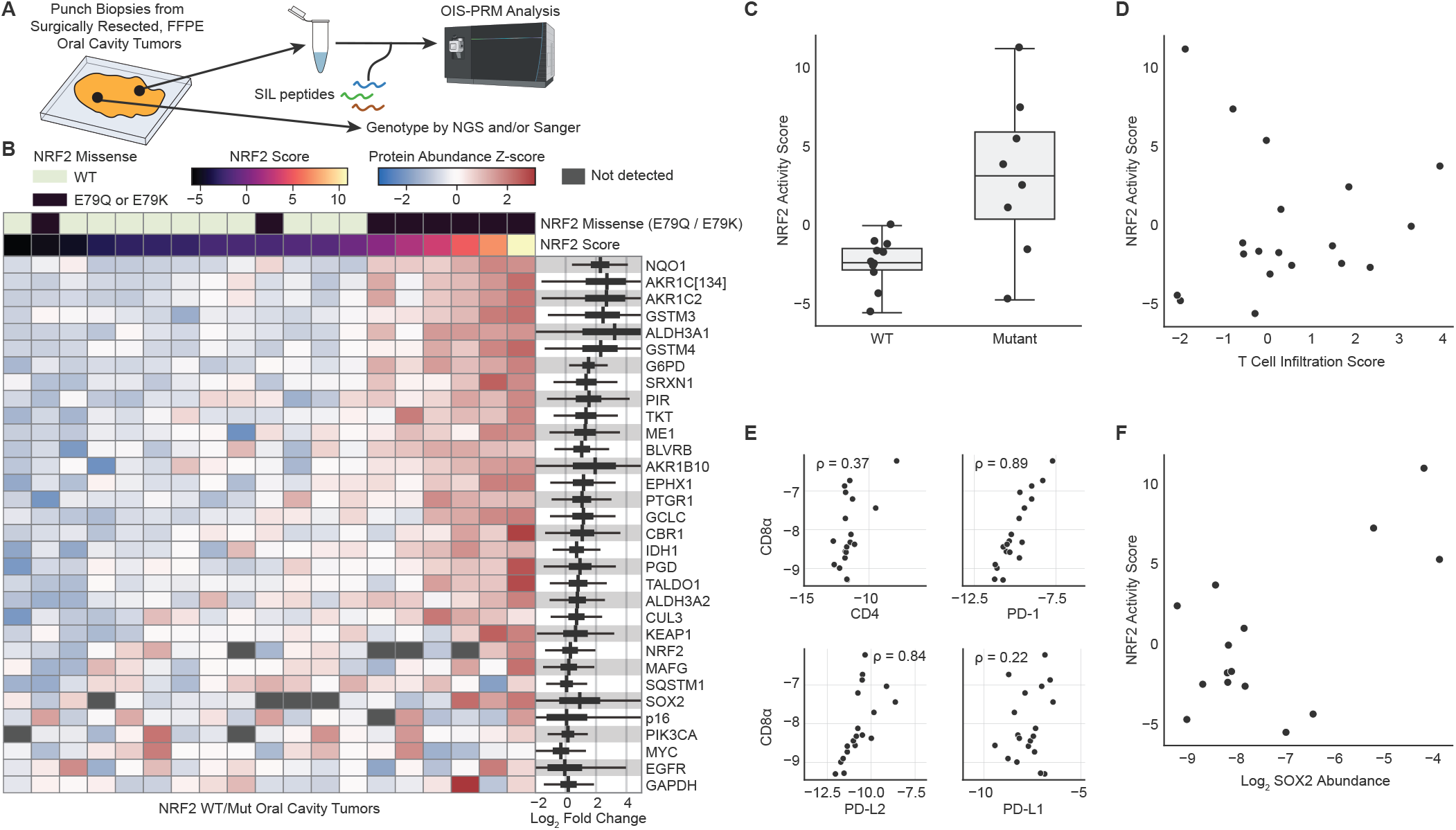
OIS-PRM analysis of oral squamous cell carcinomas showed concordance between NRF2 genotype and target gene expression. **A)** Schema describing the collection, genotyping, and OIS-PRM analysis of oral cavity tumors. Protein was extracted from punch biopsies of FFPE tumor blocks and subject to OIS-PRM. Tumors were either NRF2 WT, NRF2 E79Q, or NRF2 E79K **B)** Expression of NRF2 target proteins and others by OIS-PRM. On the right-hand-side the thick black bands contain 95% of the posterior density of the mean logarithmic fold change between the active NRF2 Mut and inactive NRF2 WT tumors. The narrow grey bands contain 95% of the posterior predictive density for the logarithmic fold change in expression between an NRF2 active over an NRF2 inactive tumor. **C)** NRF2 activity scores for the NRF2 WT and mutant (E79Q or E79K) tumors. **D)** Scatterplot of immune infiltration scores and NRF2 activity scores for each tumor. Scores were calculated by PCA as in **Figure 6** and **Figure 5** respectively. **E)** Spearman correlations between the immuno-oncology markers. **F)** Scatterplot of NRF2 activity scores and SOX2 protein abundance for each tumor.

## Discussion

This work presents an optimized targeted proteomics method called OIS-PRM and a SIL peptide library that may be valuable for basic, pre-clinical and clinical research. Within the clinical arena, biomarker assays are needed in HNSCC to predict patient response to radiation therapy, including HPV(-) patients and HPV(+) patients suffering recurrent disease. We and others recently reported that NRF2 activating genotypes predict poor response to radiation therapy, as quantified by locoregional failure following single-modality RT [13, 14]. The resulting assertion, which remains to be clinically implemented, is that HPV(-) HNSCC that are NRF2 inactive should receive standard of care radiation. Conversely, patients with NRF2-active tumors should not receive standard of care radiation dosage, but consider surgery or more aggressive therapeutic regimens. Although HPV(+) head and neck tumors rarely show high NRF2 signaling at initial presentation, recurrent HPV(+) tumors frequently harbor mutant NRF2 alleles [51]; therefore, patients with recurrent HPV(+) cancer should likewise undergo screening for high NRF2 signaling. For basic cancer research, OIS-PRM provides a powerful multiplexed protein quantitation assay that if implemented as a shared resource would be cost effective and empowering. Pre-clinical and translational sciences would also benefit from OIS-PRM. For example, though the FDA has not approved NRF2 inhibitors, future clinical trials for any such drugs could use OIS-PRM as a mechanistic biomarker.

In addition to the value of NRF2-centered biomarkers, OIS-PRM-enabled protein-level quantitation of T-cell infiltration and immune checkpoint proteins may predict patient response to anti-PD1 therapies. Current predictive biomarkers for anti-PD1 therapy response include antibody staining for PD-L1, tumor mutational burden, and an mRNA expression-based IFN-gamma signature [23, 25, 26]. OIS-PRM with an optimized SIL peptide library may prove superior to these methods, but at present the strong co-linearity between most of the immunooncology markers studied in this work limits its predictive potential. Future expansion of our SIL peptide catalogue will include additional immune checkpoint proteins, cytokines, chemokines, and markers for subtypes of innate immune cells that suppress anti-tumor immunity.

For the HPV(+) tumors in particular, lack of T-cell infiltration might predict poor overall survival independent of anti-PD1 therapy. Our work supports the finding of previous studies that HPV(+) tumors illicit stronger T-cell infiltration into the tumor microenvironment compared to HPV(-) cancers. That said, T-cell infiltration is a continuous variable across patient cohorts. A recent proteomics study reported significantly reduced expression of T-cell markers in HPV(+) tumors that would eventually recur, compared to those that did not recur [52]. Our data suggest that a small fraction of HPV(+) tumors have low T-cell infiltration comparable to that of typical HPV(-) tumors. This is an important consideration given that HPV(+) patients may receive deintensified radiation treatment regiments because of the superior survival outcomes compared to HPV(-) tumors. For patients with HPV(+) tumors that are poorly infiltrated by T cells, deintensification risks poor outcome.

With current technologies, we estimate that the upper end of an OIS-PRM assay may include 200–700 protein targets depending on gradient length and the desired sensitivity. The selection, design and construction of a SIL panel is therefore worthy of comment. Our results and analysis of publicly available CPTAC proteomics data suggest that neither genotype nor the expression of any single protein accurately predicts NRF2 pathway activity. Genotype-phenotype annotations for cancer-derived mutations remain sparse—particularly for tumor suppressor genes—and thus mutation-based classifiers often suffer high false discovery rates. It has been difficult to predict the functional effects of KEAP1 mutation on NRF2 transcriptional activity [7]. We found that protein expression NRF2 targets efficiently separated NRF2 active from NRF2 inactive tumors (Fig. 2A-C), but not all of these targets are equally diagnostic for NRF2 activity. We therefore modeled the expression of each NRF2 target in our SIL peptide library in both cell lines and oral cavity tumors to quantify to what extent and how consistently NRF2 signaling drove the expression of each target (Fig. 4B, Fig 7B). Notably, HMOX1 ranked poorly among all NRF2 targets in the panel despite its widespread use as a canonical NRF2 marker [53].

RNA biomarkers and protein biomarkers independently offer great value for personalized medicine. With the rapid technological and computational advancements in mass spectrometry, protein-based assays are nearing the comprehensiveness of genomic assays. Many features of proteins make them superior to mRNA-based biomarker assays, not the least of which are the complicated mechanisms governing the abundance of mRNA to its protein product [44, 54]. We found that for NRF2 target genes, the protein-to-mRNA correlations are moderate to strong, such that transcript abundances do well to distinguish between the NRF2 active and inactive cases (Figure 2F) [30]. However, for other proteins in our catalogue such as NRF2 itself, KEAP1, PD-L1, PD-L2, and various immune checkpoint proteins and cytokines, correlations between the mRNA and respective proteins are weak or lacking in validation [28, 30, 55].

Our OIS-PRM analysis of HNSCC tumor samples revealed varied NRF2 activity. From a small cohort of 30 HNSCC oropharynx tumors, we identified a single HPV(-) tumor with high NRF2 signaling (Fig. 5). Since approximately 14 and 5 percent of HPV(-) HNSCCs present with NRF2 and KEAP1 mutations respectively, we expected to observe at least one NRF2 active tumor [56]. Several other HPV(-) tumors presented moderately elevated NRF2 scores, perhaps owing to non-genomic mechanisms of pathway activation such as competitive KEAP1 inhibition or NRF2 copy number amplifications [7]. Whether this intermediate NRF2 activation impacts responsiveness to radiation therapy remains to be seen, but future analysis of appropriately sized training and validation cohorts could reveal a threshold of clinical relevance. Subsequent analysis of a separate cohort of NRF2 genotyped oral cavity tumors (Fig. 7) further confirmed that targeted proteomics can identify NRF2 active tumors and quantify immune checkpoint proteins and cancer drivers. However, two of the 8 tumors harboring NRF2 E79Q or E79K alleles did not overexpress NRF2 targets at the protein level. We hypothesize that spatial heterogeneity within each tumor between the genotyped punch and the independent punches taken for proteomics could explain this discrepancy.

Our data also present several unexpected observations pertaining to NRF2 driven immune-suppression, a correlation between the NRF2 and SOX2 oncogenes, and NRF2 activation in an HPV(+) background. First, given recent publications, we expected NRF2 activation to inversely correlate with T-cell infiltration [23, 32, 33]. Our data do not support this hypothesis. However, the literature evidence most strongly shows that NRF2 activity correlates with resistance to anti-PD1 drugs, drives expression of PD-L1, and supports polarization of tumor infiltrating leukocytes towards immunosuppressive functions [14, 23, 31, 35]. Therefore, it is possible that NRF2 mediates immune suppression by modulating the infiltration and function of innate immune cells rather than the abundance of T-cells at the primary tumor site. Indeed, we recently observed in mice that NRF2 activation within syngeneic grafted HNSCC tumors polarized infiltrating monocytes from an M1 towards an M2 phenotype and correlated with increased abundance of myeloid derived suppressor cells [14]. Likewise, overexpression of an NRF2 target, GPX2, in a different mouse model of oral cancer results in a strikingly similar phenotype but with a reduction in T-cell infiltration that was not observed by Guan et al [14, 33]. Clearly though, the sample size in our study is limited, thus weakening statistically meaningful observations with respect to T-cell infiltration. Secondly, Harkonen et al. recently observed correlation between SOX2 copy number and NRF2 transcriptional signature [32]. We also observed co-expression between the SOX2 and NRF2 oncogenes in head and neck cancer and believe this association merits further investigation. Finally, we observed that the second principal component of the NRF2 proteins separated HPV(+) from HPV(-) tumors, suggesting that NRF2 differently activates its target genes in an HPV(+) compared to an HPV(-) background.

Finally, several discussion points on the development of OIS-PRM are warranted. OIS-PRM differs from SureQuant^™^ and current state-of-the art methods primarily in that it efficiently orders scans within each scan cycle and monitors peptide elution in real-time to avoid acquiring uninformative scans during long peak tails. Prior art recommends rapid data acquisition to ensure capture of 6-10 data points for each peptide analyte [40, 57]. However, this heuristic rule might apply differently depending on whether quantification relies on raw peak areas or on ratios with internal standards. In theory, a single measurement should reflect the relative abundances of a SIL peptide and its endogenous counterpart, with additional scans minimizing the effects of noise and variability. TMT-labeling experiments operate on this principle and quantify peptides by the relative abundances of reporter ions in as few as one MSn scan. Accordingly, while OIS-PRM increased the number of peptides quantified with at least seven points and decreased median CVs, it failed to quantify more peptides with a CV of less than 20%. When using internal standards, however, dense chromatogram sampling enables alignment of SIL and endogenous chromatographic profiles. Poor correspondence of light and heavy counterparts reveals interfered or noisy transitions unsuitable for quantification, with the absence of aligned transitions serving as a pseudo limit of detection. Therefore, this work and a recent report describe spectral angle metrics to measure similarity between light and heavy peptides [58]. We propose that a 1-cycle delay between MS1 detection and the subsequent watch scan could explain why the SureQuant ^™^ method frequently missed peak fronts (Fig. 1I). Because of these aforementioned advantages, OIS-PRM will enable the use of even larger SIL peptide arrays, thus empowering proteomic interrogation of tumor biology and personalized medicine.

## Materials and methods

### Cell culture and lentiviral transduction

All cell lines were maintained in a humidified incubator at 37°C with 5% CO2. Cell line identities were validated by short tandem repeat analysis (LabCorp, Genetica Cell Line Testing), and cultures were regularly tested for mycoplasma contamination (Lonza). The UPCI:SCC090 (CRL-3239), UPCI:SCC152 (CRL-3240), and UPCI:SCC154 (CRL-3241) cell lines were purchased from ATCC and cultured in EMEM (Corning) supplemented with 10% FBS (Sigma), 1% penicillin–streptomycin (Corning), and 2 mmol/L L-glutamine (GIBCO). HEK293T cells (CRL-11268) were purchased from ATCC and cultured in DMEM (Corning) supplemented with 10% FBS and 1% penicillin–streptomycin.

Recombinant lentivirus was produced in HEK293T cells using polyethylenimine (PEI) based transfection. Briefly, psPAX2 packaging (Addgene #12260), VSV-G envelope (Addgene #12259), and UBC driven NRF2 E79Q vectors were combined with PEI at a 3:1 ratio (μl PEI: μg DNA). Supernatants containing virus were filtered and added to SCC90, SCC152, and SCC154 cells. Transduced cells were selected with 50 μg/mL, 50 μg/mL, and 250 μg/mL hygromycin, respectively.

### Stable isotope labeled internal standard peptides

For the HNSCC peptide library, SIL peptides were obtained in array-purity from Vivitide.

These were reconstituted and combined to a nominal abundance of 300 nM/uL per peptide. The initial library included 288 peptides. Synthetic peptides missing the expected peak in their MALDI spectra or failing detection by at least 5 transitions in MS survey runs were excluded from subsequent analyses, which left 236 peptides remaining. The Kinome SIL peptide library included 705 high-purity non-phosphorylated SIL peptides that were detectable in survey analyses. The supplementary material includes a full catalogue of these peptide libraries.

### Data-dependent acquisition LC-MS/MS

Proteins were extracted from FFPE tissues using a protocol based on Coscia et al. and Kohale et al. [46, 47]. Tryptic peptides from protein digests obtained from cell lines and tissue lysates were quantified by BCA using a peptide digest standard (Thermo Scientific, catalog no. 23225; 23295). Peptides were analyzed by an Orbitrap Eclipse Tribrid mass spectrometer (Thermo Scientific). For the cell line and FFPE tissue digests 1 ug and 1.5 ug of endogenous peptide respectively were injected per run. SIL peptides were injected at 150 fmol/peptide.

Database searching was carried out by MaxQuant version 2.0.3.0 against the UniProt human proteome (Swiss-Prot + Trembl) with an FDR of 1%. For the cell lines and FFPE tissues respectively, these were downloaded September 16, 2021 and February 18, 2023 and contained 78,120 and 79,038 sequences.

### IS-PRM

IS-PRM methods leverage SIL internal standard peptides to direct efficient acquisition of endogenous, unlabeled peptides [36]. SureQuant^™^ specifically looks for MS1 detection of expected m/z for SIL peptides once per cycle [40]. MS1 detection at sufficient intensity triggers a subsequent “watch” scan. Detection of at least five of six characteristic transitions confirms the presence of a SIL peptide. Confirmation triggers a high-resolution “quant” scan targeting the endogenous counterpart. The custom OIS-PRM algorithm implemented through the Thermo Scientific^™^ Tribrid^™^ IAPI is detailed in Supplementary Figure 2 and Supplementary Methods.

### IS-PRM data analyses

IS-PRM identification was based on the six most abundant transitions for each peptide excluding precursor, y1, y2, and b1 ions. The three most abundant transitions were used for quantification, but spectral angle contrast between light and heavy transition areas was used to exclude noisy or interfered transitions. Peak area ratios obtained by IS-PRM were normalized by global extraction from PRM (GXPRM) as proposed by Chambers et al [59]. Briefly, the intensities of commonly identified peptides that were co-isolated with the targeted peptides are used to derive a multiplicative normalization factor for each sample. Peptide abundances are summarized to the protein level by their geometric mean.

## References

1. Johnson, D.E., et al., Head and neck squamous cell carcinoma. Nat Rev Dis Primers, 2020. 6(1): p. 92.

2. Siegel, R.L., et al., Cancer statistics, 2022. CA Cancer J Clin, 2022. 72(1): p. 7–33.

3. Sabatini, M.E. and S. Chiocca, Human papillomavirus as a driver of head and neck cancers. Br J Cancer, 2020. 122(3): p. 306–314.

4. Ang, K.K., et al., Human papillomavirus and survival of patients with oropharyngeal cancer. N Engl J Med, 2010. 363(1): p. 24–35.

5. Mody, M.D., et al., Head and neck cancer. Lancet, 2021. 398(10318): p. 2289–2299.

6. Mehra, R., et al., Efficacy and safety of pembrolizumab in recurrent/metastatic head and neck squamous cell carcinoma: pooled analyses after long-term follow-up in KEYNOTE-012. Br J Cancer, 2018. 119(2): p. 153–159.

7. Cloer, E.W., et al., NRF2 Activation in Cancer: From DNA to Protein. Cancer Res, 2019. 79(5): p. 889–898.

8. Shibata, T., et al., Cancer related mutations in NRF2 impair its recognition by Keap1-Cul3 E3 ligase and promote malignancy. Proc Natl Acad Sci U S A, 2008. 105(36): p. 13568–73.

9. Singh, A., et al., Gain of Nrf2 function in non-small-cell lung cancer cells confers radioresistance. Antioxid Redox Signal, 2010. 13(11): p. 1627–37.

10. Namani, A., et al., Gene-expression signature regulated by the KEAP1-NRF2-CUL3 axis is associated with a poor prognosis in head and neck squamous cell cancer. BMC Cancer, 2018. 18(1): p. 46.

11. Noh, J.K., et al., SOD2- and NRF2-associated Gene Signature to Predict Radioresistance in Head and Neck Cancer. Cancer Genomics Proteomics, 2021. 18(5): p. 675–684.

12. Matsuoka, Y., et al., The antioxidative stress regulator Nrf2 potentiates radioresistance of oral squamous cell carcinoma accompanied with metabolic modulation. Lab Invest, 2022. 102(8): p. 896–907.

13. Sheth, S., et al., Correlation of alterations in the KEAP1/CUL3/NFE2L2 pathway with radiation failure in larynx squamous cell carcinoma. Laryngoscope Investig Otolaryngol, 2021. 6(4): p. 699–707.

14. Guan, L., et al., NFE2L2 mutations enhance radioresistance in head and neck cancer by modulating intratumoral myeloid cells. Cancer Res, 2023.

15. Gavrielatou, N., et al., Biomarkers for immunotherapy response in head and neck cancer. Cancer Treat Rev, 2020. 84: p. 101977.

16. Cullinan, S.B., et al., The Keap1-BTB protein is an adaptor that bridges Nrf2 to a Cul3-based E3 ligase: oxidative stress sensing by a Cul3-Keap1 ligase. Mol Cell Biol, 2004. 24(19): p. 8477–86.

17. Kobayashi, A., et al., Oxidative stress sensor Keap1 functions as an adaptor for Cul3-based E3 ligase to regulate proteasomal degradation of Nrf2. Mol Cell Biol, 2004. 24(16): p. 7130–9.

18. Zhang, D.D., et al., Keap1 is a redox-regulated substrate adaptor protein for a Cul3-dependent ubiquitin ligase complex. Mol Cell Biol, 2004. 24(24): p. 10941–53.

19. Yamamoto, M., T.W. Kensler, and H. Motohashi, The KEAP1-NRF2 System: a Thiol-Based Sensor-Effector Apparatus for Maintaining Redox Homeostasis. Physiol Rev, 2018. 98(3): p. 1169–1203.

20. Cuadrado, A., et al., Therapeutic targeting of the NRF2 and KEAP1 partnership in chronic diseases. Nat Rev Drug Discov, 2019. 18(4): p. 295–317.

21. Yagishita, Y., et al., Current Landscape of NRF2 Biomarkers in Clinical Trials. Antioxidants (Basel), 2020. 9(8).

22. Forster, M.D. and M.J. Devlin, Immune Checkpoint Inhibition in Head and Neck Cancer. Front Oncol, 2018. 8: p. 310.

23. Cristescu, R., et al., Pan-tumor genomic biomarkers for PD-1 checkpoint blockade-based immunotherapy. Science, 2018. 362(6411).

24. Grossman, J.E., et al., Is PD-L1 a consistent biomarker for anti-PD-1 therapy? The model of balstilimab in a virally-driven tumor. Oncogene, 2021. 40(8): p. 1393–1395.

25. Haddad, R.I., et al., Influence of tumor mutational burden, inflammatory gene expression profile, and PD-L1 expression on response to pembrolizumab in head and neck squamous cell carcinoma. J Immunother Cancer, 2022. 10(2).

26. Ayers, M., et al., IFN-gamma-related mRNA profile predicts clinical response to PD-1 blockade. J Clin Invest, 2017. 127(8): p. 2930–2940.

27. Morales-Betanzos, C.A., et al., Quantitative Mass Spectrometry Analysis of PD-L1 Protein Expression, N-glycosylation and Expression Stoichiometry with PD-1 and PD-L2 in Human Melanoma. Mol Cell Proteomics, 2017. 16(10): p. 1705–1717.

28. Liebler, D.C., et al., Analysis of Immune Checkpoint Drug Targets and Tumor Proteotypes in Non-Small Cell Lung Cancer. Sci Rep, 2020. 10(1): p. 9805.

29. Wang, D., et al., A deep proteome and transcriptome abundance atlas of 29 healthy human tissues. Molecular Systems Biology, 2019. 15(2): p. e8503.

30. Nusinow, D.P., et al., Quantitative Proteomics of the Cancer Cell Line Encyclopedia. Cell, 2020. 180(2): p. 387–402.e16.

31. Zhu, B., et al., Targeting the upstream transcriptional activator of PD-L1 as an alternative strategy in melanoma therapy. Oncogene, 2018. 37(36): p. 4941–4954.

32. Harkonen, J., et al., A pan-cancer analysis shows immunoevasive characteristics in NRF2 hyperactive squamous malignancies. Redox Biol, 2023. 61: p. 102644.

33. Ahmed, K.M., et al., Glutathione peroxidase 2 is a metabolic driver of the tumor immune microenvironment and immune checkpoint inhibitor response. J Immunother Cancer, 2022. 10(8).

34. Papillon-Cavanagh, S., et al., STK11 and KEAP1 mutations as prognostic biomarkers in an observational real-world lung adenocarcinoma cohort. ESMO Open, 2020. 5(2).

35. Papalexi, E., et al., Characterizing the molecular regulation of inhibitory immune checkpoints with multimodal single-cell screens. Nat Genet, 2021. 53(3): p. 322–331.

36. Gallien, S., S.Y. Kim, and B. Domon, Large-Scale Targeted Proteomics Using Internal Standard Triggered-Parallel Reaction Monitoring (IS-PRM). Mol Cell Proteomics, 2015. 14(6): p. 1630–44.

37. Stopfer, L.E., et al., High-Density, Targeted Monitoring of Tyrosine Phosphorylation Reveals Activated Signaling Networks in Human Tumors. Cancer Res, 2021. 81(9): p. 2495–2509.

38. Stopfer, L.E., et al., Absolute quantification of tumor antigens using embedded MHC-I isotopologue calibrants. Proc Natl Acad Sci U S A, 2021. 118(37).

39. Nguyen, C.D.L., et al., A sensitive and simple targeted proteomics approach to quantify transcription factor and membrane proteins of the unfolded protein response pathway in glioblastoma cells. Sci Rep, 2019. 9(1): p. 8836.

40. Gajadhar, A., SureQuant intelligence-driven MS: a new paradigm for targeted quantitation. 2020, Thermo Fisher Scientific.

41. Satpathy, S., et al., A proteogenomic portrait of lung squamous cell carcinoma. Cell, 2021. 184(16): p. 4348–4371 e40.

42. Huang, C., et al., Proteogenomic insights into the biology and treatment of HPV-negative head and neck squamous cell carcinoma. Cancer Cell, 2021. 39(3): p. 361–379 e16.

43. Gillette, M.A., et al., Proteogenomic Characterization Reveals Therapeutic Vulnerabilities in Lung Adenocarcinoma. Cell, 2020. 182(1): p. 200–225 e35.

44. Liu, Y., A. Beyer, and R. Aebersold, On the Dependency of Cellular Protein Levels on mRNA Abundance. Cell, 2016. 165(3): p. 535–50.

45. Bowman, B.M., et al., A conditional mouse expressing an activating mutation in NRF2 displays hyperplasia of the upper gastrointestinal tract and decreased white adipose tissue. J Pathol, 2020. 252(2): p. 125–137.

46. Coscia, F., et al., A streamlined mass spectrometry–based proteomics workflow for large-scale FFPE tissue analysis. The Journal of Pathology, 2020. 251(1): p. 100–112.

47. Kohale, I.N., et al., Quantitative Analysis of Tyrosine Phosphorylation from FFPE Tissues Reveals Patient-Specific Signaling Networks. Cancer Research, 2021. 81(14): p. 3930–3941.

48. Marchione, D.M., et al., HYPERsol: High-Quality Data from Archival FFPE Tissue for Clinical Proteomics. Journal of Proteome Research, 2020. 19(2): p. 973–983.

49. Barnabas, G.D., et al., ASAP horizontal line Automated Sonication-Free Acid-Assisted Proteomes horizontal line from Cells and FFPE Tissues. Anal Chem, 2023.

50. Hong, S., et al., Topoisomerase IIbeta-binding protein 1 activates expression of E2F1 and p73 in HPV-positive cells for genome amplification upon epithelial differentiation. Oncogene, 2019. 38(17): p. 3274–3287.

51. Vyas, A., et al., Recurrent Human Papillomavirus-Related Head and Neck Cancer Undergoes Metabolic Reprogramming and Is Driven by Oxidative Phosphorylation. Clin Cancer Res, 2021. 27(22): p. 6250–6264.

52. Kaneko, T., et al., Proteome and phosphoproteome signatures of recurrence for HPV(+) head and neck squamous cell carcinoma. Commun Med (Lond), 2022. 2: p. 95.

53. Levings, D.C., et al., A distinct class of antioxidant response elements is consistently activated in tumors with NRF2 mutations. Redox Biol, 2018. 19: p. 235–249.

54. Goncalves, E., et al., Widespread Post-transcriptional Attenuation of Genomic Copy-Number Variation in Cancer. Cell Syst, 2017. 5(4): p. 386–398 e4.

55. Israelsson, P., et al., Cytokine mRNA and protein expression by cell cultures of epithelial ovarian cancer-Methodological considerations on the choice of analytical method for cytokine analyses. Am J Reprod Immunol, 2020. 84(1): p. e13249.

56. Cancer Genome Atlas, N., Comprehensive genomic characterization of head and neck squamous cell carcinomas. Nature, 2015. 517(7536): p. 576–82.

57. Gallien, S., et al., Technical considerations for large-scale parallel reaction monitoring analysis. J Proteomics, 2014. 100: p. 147–59.

58. Bourmaud, A., S. Gallien, and B. Domon, Parallel reaction monitoring using quadrupole-Orbitrap mass spectrometer: Principle and applications. Proteomics, 2016. 16(15-16): p. 2146–59.

59. Chambers, A.G., et al., Global Extraction from Parallel Reaction Monitoring to Quantify Background Peptides for Improved Normalization and Quality Control in Targeted Proteomics. Analytical Chemistry, 2021. 93(40): p. 13434–13440.

