## Supplemental Figures for "Targeted proteomic quantitation of NRF2 signaling and predictive biomarkers in HNSCC": SF1.pdf

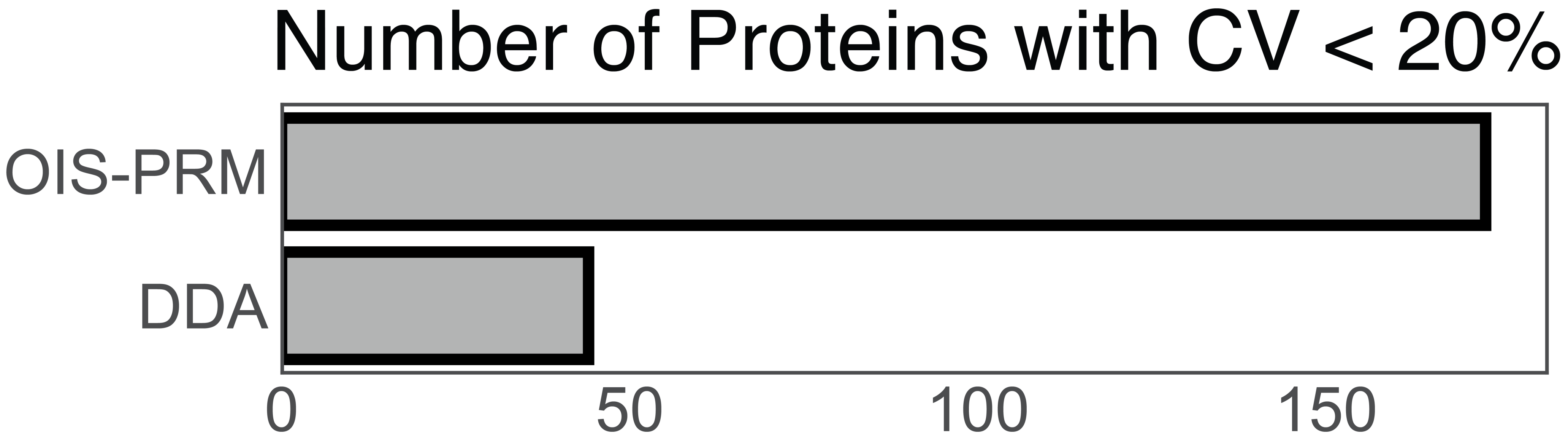


**OIS-PRM Improves Sensitivity over DDA**. OIS-PRM quantifies more proteins than does DDA (172 vs. 47) a with a coefficient of variation (CV) less than 20%. See main figure 1.
