## Supplemental Figures for "Targeted proteomic quantitation of NRF2 signaling and predictive biomarkers in HNSCC": SF2.pdf

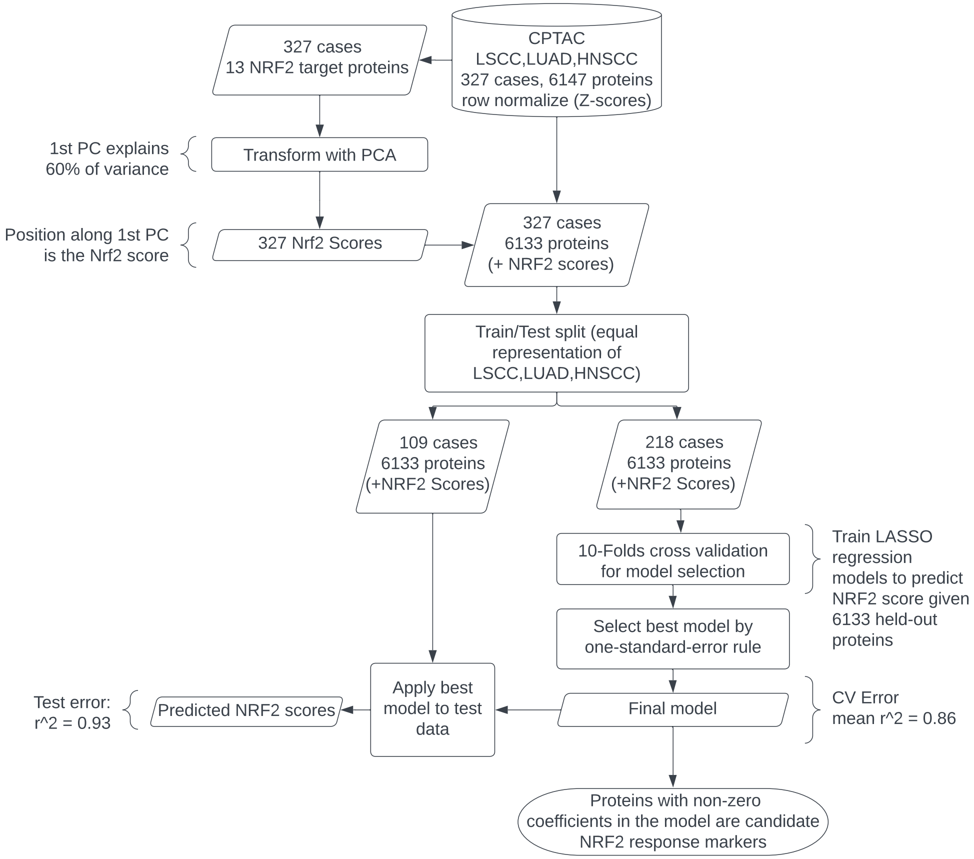


**CPTAC Analysis.** Schema of NRF2 pathway analysis from CPTAC cohorts for HNSCC, LUSC, and LUAD. See main figure 2.
