## Supplemental Figures for "Targeted proteomic quantitation of NRF2 signaling and predictive biomarkers in HNSCC": SF3.pdf

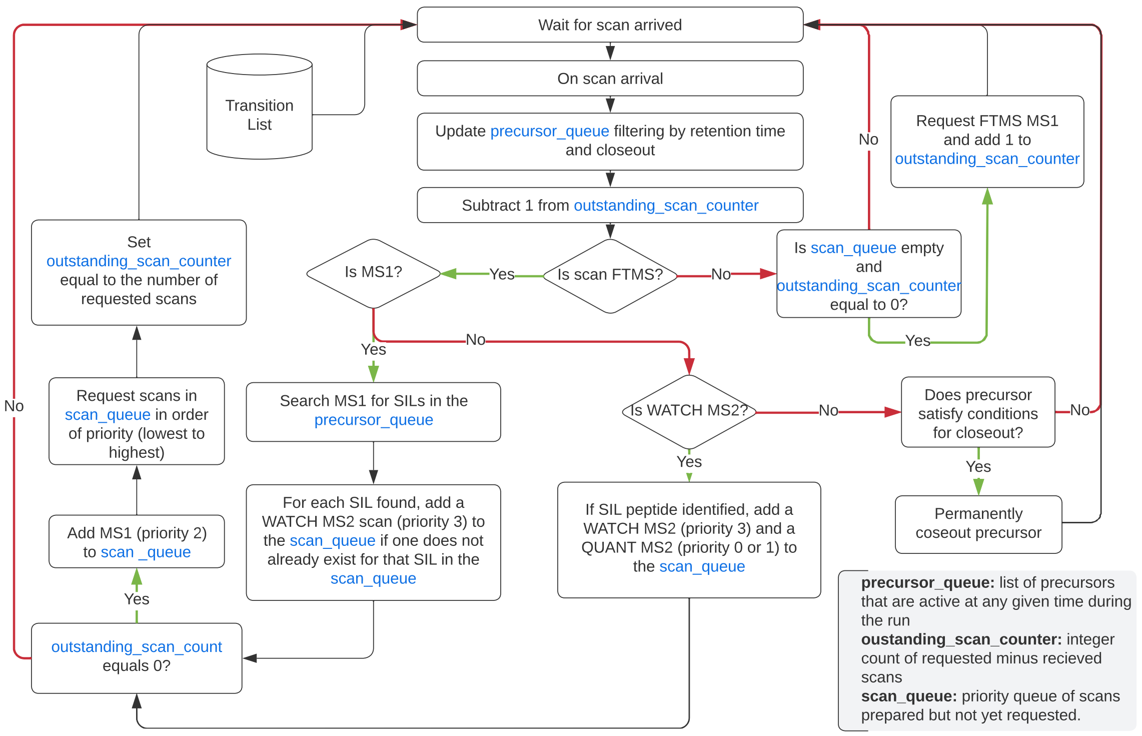


**OIS-PRM Algorithm**. Schematic of the OIS-PRM data acquisition algorithm as detailed in supplemental methods.
