## Supplemental Figures for "Targeted proteomic quantitation of NRF2 signaling and predictive biomarkers in HNSCC": SF4.pdf

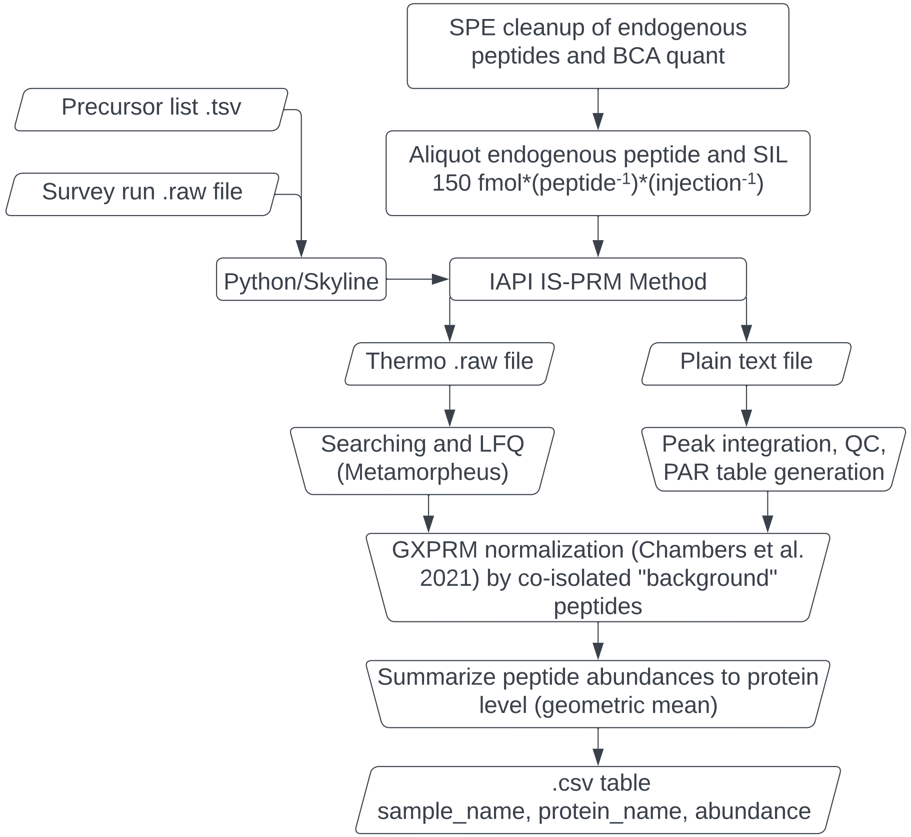


**OIS-PRM Analysis Pipeline.** Schematic of the data analysis pipeline for OIS-PRM and SureQuant™ experiments as detailed in supplemental methods.
