## Supplemental Figures for "Targeted proteomic quantitation of NRF2 signaling and predictive biomarkers in HNSCC": SF5.pdf

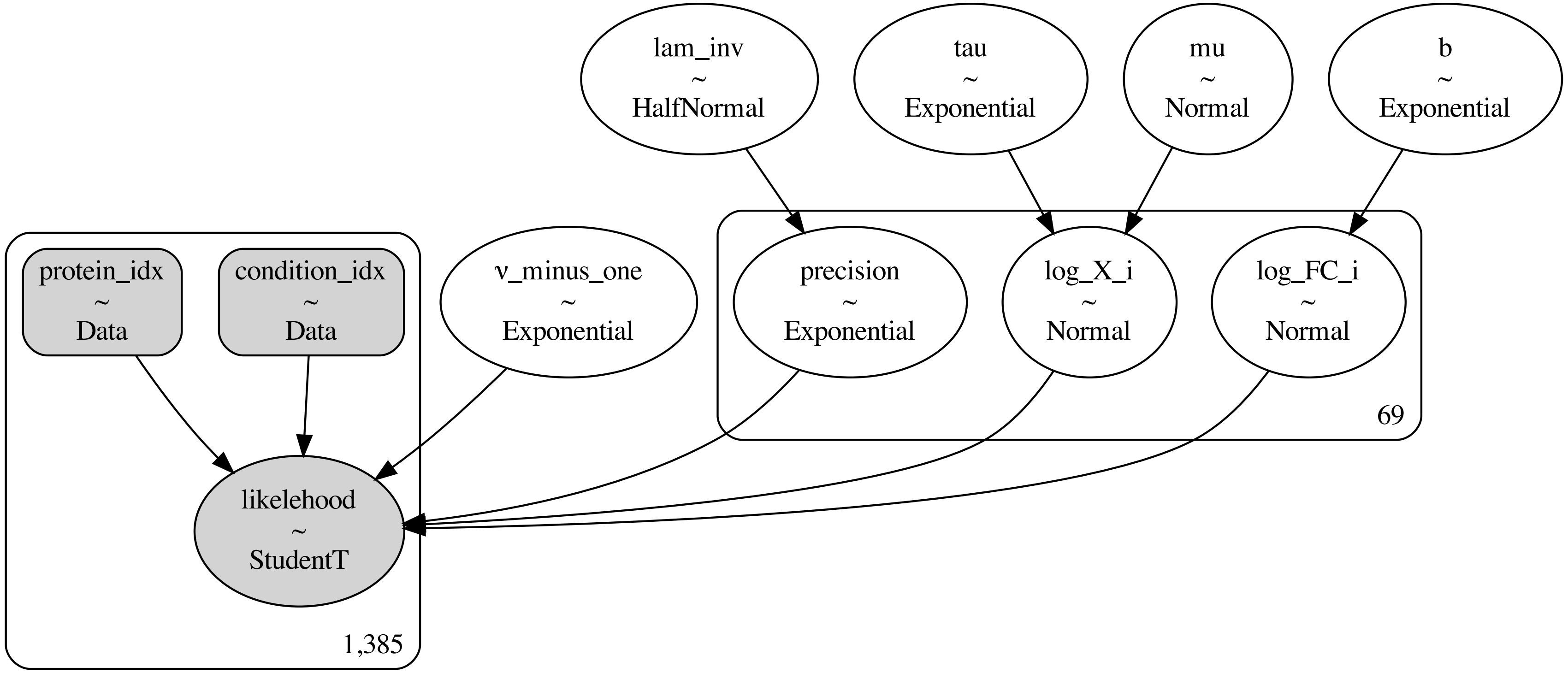


**NRF2 Target Expression Model.** Plate diagram for hierarchical Bayesian model used to estimate posterior distributions for mean fold changes in the expression of NRF2 targets between NRF2 active and inactive cell lines and tumors. Detailed in supplemental methods.
